# Optical Fibers Functionalized with Single-Walled Carbon Nanotubes for Flexible Fluorescent Catecholamine Detection

**DOI:** 10.1101/2024.07.09.602792

**Authors:** Madeline E. Klinger, Rigney A. Miller, Linda Wilbrecht, Markita P. Landry

## Abstract

Despite the blockbuster popularity of drugs that act on catecholamine receptors, catecholamine dynamics in human health and disease remain an incomplete picture. Recent advances in fluorescent sensors have enabled unprecedented access to catecholamine dynamics in preclinical animal models, but the requirements of these technologies limit translational value for clinical diagnostics. Here, we present a flexible and convenient tool for fluorescent catecholamine detection by functionalizing optical fibers with single-walled carbon nanotube (SWNT)-based near-infrared catecholamine sensors (nIRCats), a form factor that has potential for more convenient and less invasive clinical translation. We show that these near-infrared functionalized (nIRF) fibers respond to dopamine in a biologically-relevant concentration range (10nM through 1 μM) with a mean Δ*F*/*F*_*0*_ of 0.022 through 0.411, with no statistically significant effect on signal magnitude after 16-hour exposure to human blood plasma. We further demonstrate the utility of these fibers in as little as 10 μL volumes of clinically relevant biofluids up to 24 weeks after preparation, with a Δ*F*/*F*_*0*_ of up to 0.059 through 1.127 for 10 nM through 1 μM dopamine. We also introduce a compact, mobile dual-near-infrared fiber photometry rig and demonstrate its success detecting dopamine with 0.005 Δ*F*/*F*_*0*_ in acute brain slices with nIRF fibers. Together, this fiber-based tool and photometry rig expand the toolbox of catecholamine detection technologies to a broader range of applications.

---

The catecholamine dopamine has been implicated in movement and reward prediction^1–3^, neurodegenerative diseases such as Parkinson’s and Alzheimer’s Diseases^4–7^, neuropsychiatric disorders^8–10^, and physiological functions^11–16^. Dopamine is hydroxylated to norepinephrine (noradrenaline), which is similarly implicated in neurological and psychiatric illness^17–19^, stress and arousal^20–23^, memory^24–26^, and other functions^27^. While the last few decades of research have demonstrated interactions between dopamine and norepinephrine in health and disease^17,18,28–35^, our understanding is far from complete, and physicians have limited tools to monitor catecholamine levels in human patients^36^. To accelerate this field of study, we need versatile technologies to examine catecholamine dynamics in settings from preclinical animal models to human patients.

In recent years, non-genetically encoded fluorescent molecules have emerged as viable tools for bioimaging both *ex vivo* and *in vivo*. In particular, single-walled carbon nanotubes (SWNTs) have demonstrated remarkable flexibility as biocompatible biosensors, capable of detecting DNA polymorphisms, nitric oxide, proteins, and other biomarkers^37–39^. These sensors rely on the intrinsic near-infrared (nIR) fluorescence of SWNTs, which can be modulated by adsorbed polymers to emit only in the presence of select analytes. Functionalized SWNTs emit at 1000-1300 nm, well within the nIR-II “second window” of optimal biological imaging, in which tissue absorbance and scattering of photons is minimal^40^. Functionalized SWNTs employed in drug delivery investigations have demonstrated low toxicity *in vivo*^41^. Examination of blood serum biomarkers post-intravenous injection of DNA-functionalized SWNTs suggests long-term (∼5 months) biocompatibility in rodent models^42^, and implanted SWNT-based chemical sensors have previously been used to measure nitric oxide in mice for over 400 days^43^. These features position SWNT-based biosensors as a promising candidate for long-term biological *ex vivo* and *in vivo* imaging, where their synthetic nature facilitates easy adoption for use across animal species.

Here, we leverage the functional properties of SWNTs to develop optical fibers surface-functionalized with near-infrared catecholamine nanosensors (nIRCats) to enable rapid detection of catecholamines in small volumes of biofluids without contamination of samples. nIRCats are SWNTs functionalized with (GT)_6_ single-stranded DNA that emit up to a 24-fold and 35-fold increase in fluorescence (Δ*F*/*F*_*0*_) in response to dopamine and norepinephrine, respectively, *in vitro*^44^. To date, we have validated the use of nIRCats in solution for measuring evoked dopamine release in *ex vivo* striatal brain tissue with micron-level spatial resolution and millisecond-level temporal resolution, in the presence of pharmacological agents, and in evaluating Huntington’s Disease progression^44,45^. nIRCats have also been used on immobilized glass surfaces to measure neuronal dopamine signaling on a sub-cellular scale^46,47^. Immobilizing nIRCats on optical fibers couples our catecholamine-sensing technology with fiber photometry, a commonly employed method in systems and behavioral neuroscience, thus expanding the available imaging toolkit to permit a more adaptable experimental paradigm for a wider variety of applications. Using these nIRCat-functionalized optical (nIRF) fibers, we demonstrate a fluorescent response to 10 nM dopamine in as little as 10 μL biofluid *in vitro* and record endogenous dopamine transients evoked by electrical stimulation in *ex vivo* mouse brain tissue. We show that nIRF fibers are shelf-stable for 24 weeks post-synthesis, reusable, and non-biofouling in human plasma. Together, these results suggest translatable potential for real-time readout of catecholamine release in biological environments.

We developed a protocol for aminosilane-based immobilization^48^ of nIRCats on glass optical fibers. 400 μm diameter multimode silica optical fibers were cleaned, then silanized with 5% (3-Aminopropyl)triethoxysilane (APTES) solution in toluene. After overnight curing at 110°C, fibers were passively incubated in nIRCat solution to promote an electrostatic interaction between the DNA adsorbed on the SWNT backbone and the aminosilane bound to the silica of the fiber (**Fig. 1A**). We first assessed the fluorescent spectra of these nIRCat-functionalized (nIRF) fibers in a 40 mL solution of 1x phosphate buffered saline (PBS) to approximate saline conditions of biological environments. In agreement with the known dopamine response of solution-phase nIRCat^44,49^, we observed a response profile with characteristic peaks at 1150 and 1200 nm from all fibers. To assess the functionality of immobilized nIRCats, we added aliquots of concentrated dopamine solution directly to the PBS solution to produce sequential solutions of 10 nM, 100 nM, and 1 μM dopamine. As with solution-phase nIRCat, we observed a stepwise increase in fluorescent response above baseline fluorescence, Δ*F*/*F*_*0*_, to each subsequent addition of dopamine in 16/21 fibers, though the individual response magnitude varied between replicates (10 nM dopamine = 0.022 ± 0.056 Δ*F*/*F*_*0*_, 100 nM dopamine = 0.223 ± 0.162 Δ*F*/*F*_*0*_, and 1μM dopamine = 0.411 ± 0.230 Δ*F*/*F*_*0*_ (means ± SD); *n* = 16; *P* < 0.001 between 10 nM versus 100 nM dopamine and *P* = 0.015 between 100 nM versus 1μM dopamine, t-test; **Fig. 1B,C**). The remaining 5 fibers exhibited a decrease in Δ*F*/*F*_*0*_ to 1 μM relative to 100 nM dopamine and/or yielded Δ*F*/*F*_*0*_ < 0.1 to 1 μM dopamine and thus were deemed non-functional and excluded from further analysis (Fig. S1).

**Figure 1.**
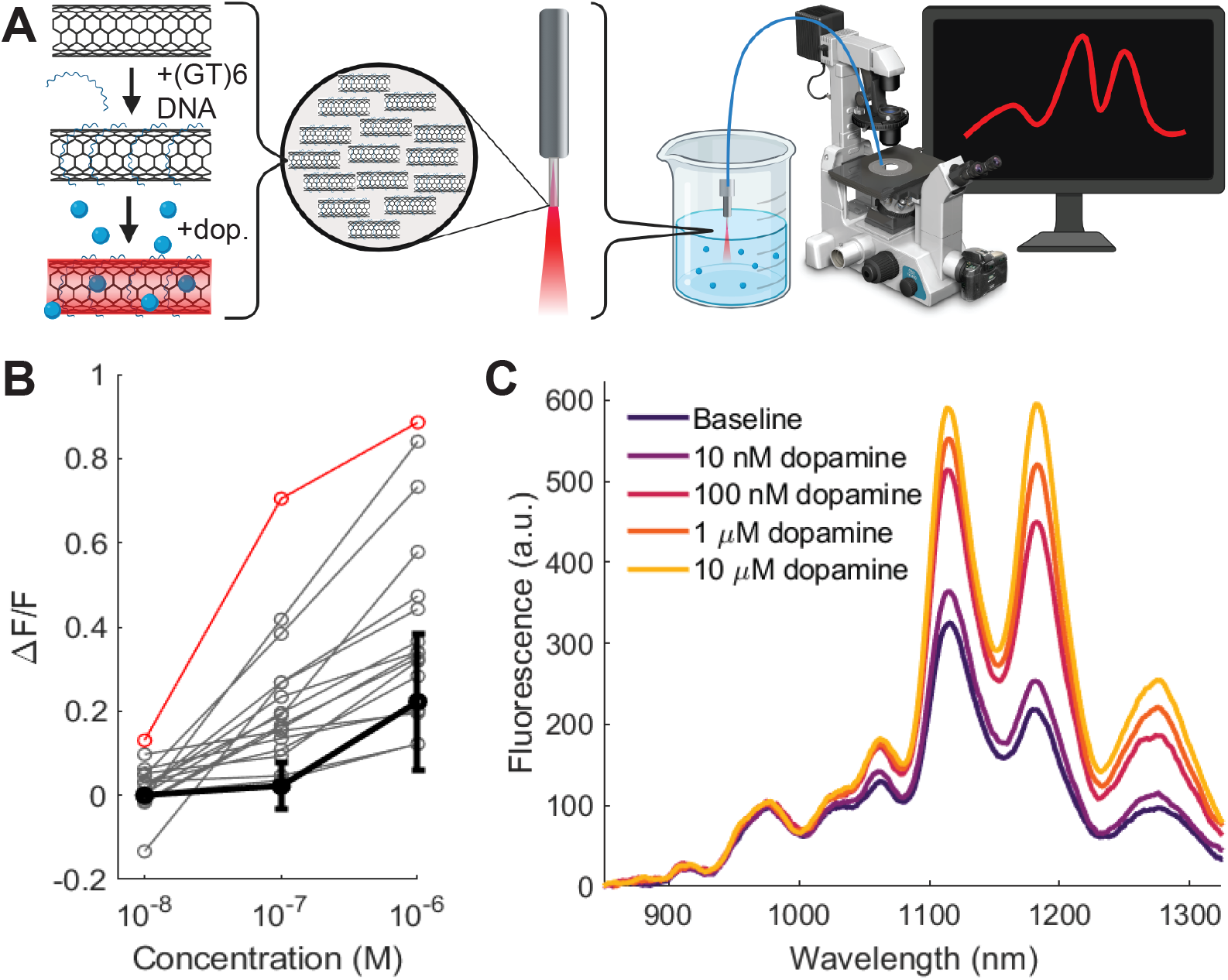
**A)** Schematic of fiber development. **B)** Integrated fluorescence from 1050-1300nm for increasing concentrations of dopamine for individual fibers (gray lines, *n* = 16) and mean (black line; error bars = SD). **C)** Representative spectra from one fiber (red in panel B) in 1x PBS and in increasing concentrations of dopamine *in vitro*.

To assess the impact of potential biofouling by molecules present in biological environments^50^, we incubated fibers (*n* = 6) in 10x diluted human blood plasma overnight (>16 hrs) and found no effect on dopamine response the next day (before incubation: 10 nM dopamine = 0.036 ± 0.050 Δ*F*/*F*_*0*_, 100 nM dopamine = 0.271 ± 0.202 Δ*F*/*F*_*0*_, and 1 μM dopamine = 0.390 ± 0.242 Δ*F*/*F*; after incubation: 10 nM dopamine = 0.033 ± 0.025 Δ*F*/*F*_*0*_, 100 nM dopamine = 0.198 ± 0.118 Δ*F*/*F*, and 1 μM dopamine = 0.381 ± 0.189 Δ*F*/*F*_*0*_; *P* = 0.623 between time points, repeated-measures ANOVA; **Fig. 2A**). As blood plasma contains only a small subset of biomolecules present in the living brain, we next incubated a new batch of fibers (*n* = 4) in brain homogenate solution for 48 hours and observed a noticeable, but not statistically significant, attenuation of response (before incubation: 10 nM dopamine = 0.080 ± 0.099 Δ*F*/*F*_*0*_, 100 nM dopamine = 0.308 ± 0.156 Δ*F*/*F*_*0*_, and 1 μM dopamine = 0.536 ± 0.306 Δ*F*/*F*_*0*_; after incubation: 10 nM dopamine = -0.004 ± 0.053 Δ*F*/*F*_*0*_, 100 nM dopamine = 0.098 ± 0.050 Δ*F*/*F*_*0*_, and 1 μM dopamine = 0.278 ± 0.073 Δ*F*/*F*_*0*_; *P* = 0.222 between time points, repeated-measures ANOVA; **Fig. 2B**), indicating that nIRF fibers may perform successfully in biological environments.

**Figure 2.**
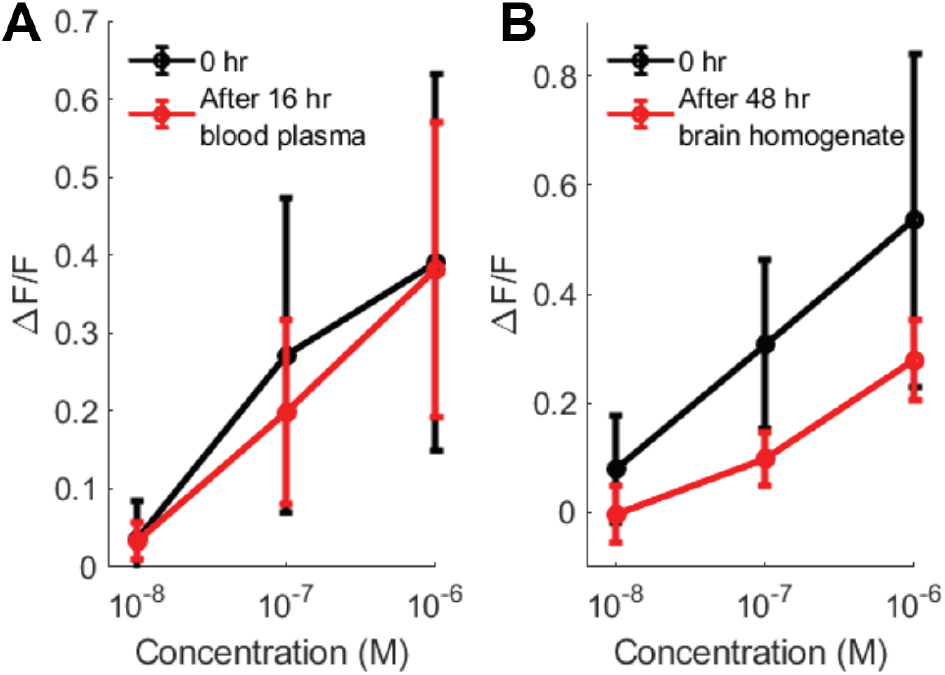
Average nIRF fluorescent responses to increasing concentrations of dopamine (error bars = SD; n = 6). A) Before plasma incubation (black) vs. after overnight plasma incubation (red). B) Before brain homogenate incubation (black) vs. after 48 hour brain homogenate incubation (red).

We next sought to study nIRF fiber extended storage time and use in a broader range of biological environments. We prepared a new batch of fibers (*n* = 4) for long-term dry storage. After 24 weeks, despite a nearly 50% reduction in baseline fluorescence, nIRF fibers could reliably detect dopamine without a statistically significant attenuation in fluorescent response relative to the initial performance (date of manufacture: 10 nM dopamine = 0.002 ± 0.020 Δ*F*/*F*_*0*_, 100 nM dopamine = 0.414 ± 0.378 Δ*F*/*F*_*0*_, and 1μM dopamine = 0.687 ± 0.355 Δ*F*/*F*_*0*_; after 24 weeks: 10 nM dopamine = 0.024 ± 0.053 Δ*F*/*F*_*0*_, 100 nM dopamine = 0.266 ± 0.174 Δ*F*/*F*_*0*_, and 1 μM dopamine = 0.517 ± 0.235 Δ*F*/*F*_*0*_; *P* = 0.479 between time points, repeated-measures ANOVA; **Fig. 3A, B**). These data suggest nIRF fibers are shelf-stable, reusable, and generate repeatable results, ideal for experimental or clinical applications.

**Figure 3.**
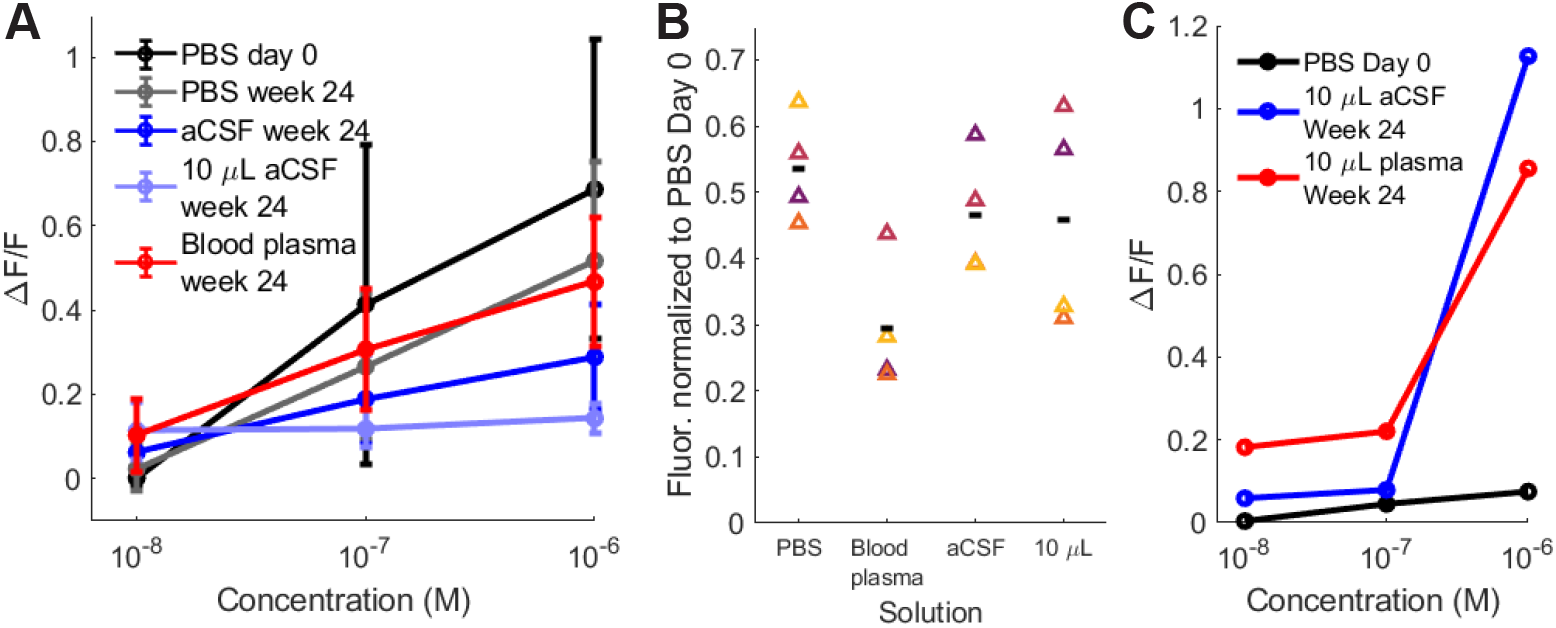
**A)** Average nIRF fiber response in biologically relevant solutions (*n* = 4; error bars = SD) **B)** Baseline fluorescence after 24 weeks in tested solutions relative to baseline fluorescence in 1x PBS on manufacturing date (colored triangles = individual fibers, black bar = mean) **C)** Single fiber response profile in clinically relevant solution volumes.

We next sought to explore the utility of nIRF fibers in another potential clinical application: biofluid samples commonly used in diagnostic testing. We next tested these same 24-week-old fibers in artificial cerebrospinal fluid (aCSF) and 100x diluted human blood plasma. While heretofore, tests had been performed in 40 mL solution, we modified our protocol to assess nIRF fiber performance in as little as 10 μL volume to account for the reduced clinical availability of patient biofluids. In all biofluid solutions, though baseline fluorescence was attenuated compared to the baseline in 1x PBS on the manufacturing date, responses to 10 nM dopamine were comparable to 10 nM dopamine in 1x PBS (40 mL 100x diluted blood plasma: 10 nM dopamine = 0.103 ± 0.086 Δ*F*/*F*_*0*_, 100 nM dopamine = 0.307 ± 0.143 Δ*F*/*F*_*0*_, and 1 μM dopamine = 0.467 ± 0.152 Δ*F*/*F*_*0*_; 40 mL aCSF: 10 nM dopamine = 0.063 ± 0.048 Δ*F*/*F*_*0*_, 100 nM dopamine = 0.190 ± 0.106 Δ*F*/*F*_*0*_, and 1 μM dopamine = 0.289 ± 0.126 Δ*F*/*F*_*0*_; 10 μL aCSF: 10 nM dopamine = 0.114 ± 0.069 Δ*F*/*F*_*0*_, 100 nM dopamine = 0.118 ± 0.043 Δ*F*/*F*_*0*_, and 1 μM dopamine = 0.143 ± 0.035 Δ*F*/*F*_*0*_; 40 mL 1x PBS listed above; *P* = 0.872 between 40 mL 1x PBS versus 40 mL blood plasma, *P* = 0.344 between 40 mL 1x PBS versus 40 mL aCSF, *P* = 0.394 between 40 mL aCSF versus 10 μL aCSF, all repeated-measures ANOVA; **Fig. 3A, B, Table S1**). As proof of principle, we further assessed the response of one fiber in 10 μL undiluted blood plasma. Compared to initial performance in PBS, this fiber exhibited a greater response to dopamine in undiluted human blood plasma after 24 weeks, with 0.182 Δ*F*/*F*_*0*_, 0.221 Δ*F*/*F*_*0*_, and 0.856 Δ*F*/*F*_*0*_ to 10 nM, 100 nM, and 1μM dopamine, respectively (**Fig. 3C**). Taken together, these data show that nIRF fibers can detect dopamine in small volumes of human biofluids, supporting the clinical utility of nIRF fibers in indicating potential risk factors for complex human disease.

We next targeted detection of evoked dopamine release from *ex vivo* dorsal striatum, an area of the brain densely innervated by dopaminergic fibers. While endogenous dopamine release in the dorsal striatum in response to reward in the intact brain ranges from about 1-20 nM^51^, electrically evoked dopamine in the same region can extend into the micromolar range^52^, well within the detection range of nIRF fibers. We constructed a compact, mobile dual-nIR fiber photometry rig with a 635nm excitation laser, ultrasensitive thermoregulated nIR camera (Ninox 640, Raptor Photonics), and flexible patch cord (**Fig. 4A**) and prepared 300 μm-thick slices of mouse striatum in a bath continual perfused with aCSF, as in previous work with solution-phase nIRCats^44,45^. We implanted a platinum iridium stimulating electrode and a nIRF fiber in the same slice and recorded a small but consistent increase in fluorescence corresponding to stimulation over four consecutive trials (mean peak Δ*F*/*F*_*0*_ = 0.005), providing evidence of endogenous dopamine release from *ex vivo* brain tissue (**Fig. 4B**). This experiment further serves as a proof of principle that, despite attenuation of signal after exposure to brain tissue, dopamine detection within a biological environment is feasible with nIRF fibers.

**Figure 4.**
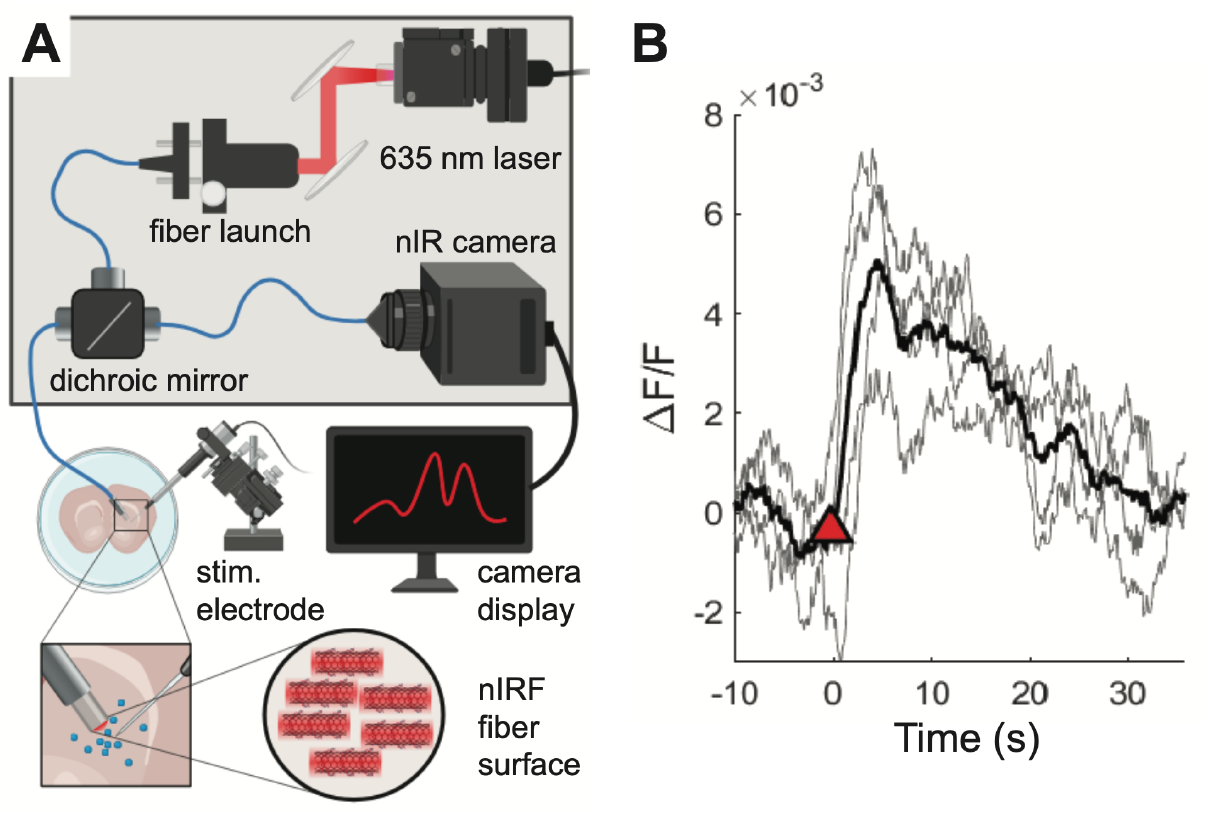
A) Schematic of 12” x 18” breadboard-based fiber photometry rig compatible with other recording platforms. B) Baseline-corrected fluorescent response to electrically evoked DA release in excised brain tissue. Gray traces correspond to individual trials (*n* = 4, one fiber); black = average; red triangle indicates time of stimulation.

While we observed consistent responses from most nIRF fibers, we also observed that some fibers were non-responsive to dopamine, which we attributed to the variable availability or accessibility of silanol groups for nIRCat bonding on the fiber surface. We thus tested protocol modifications (summarized in **Table S2**) toward the goal of regularizing fiber output. To promote greater availability of silanol groups, we treated fibers with a base bath of potassium hydroxide in ethanol and then hydroxylated the silica surface in a solution of concentrated sulfuric acid and 30% hydrogen peroxide (piranha solution). Finally, we bath sonicated fibers in nIRCat solution to facilitate interaction between nIRCats and the fiber surface. These protocol modifications increased the average baseline fluorescent response nearly 4-fold relative to the originally described protocol (*P* = 0.0012, Wilcoxon ranked sum test); this greater density of bound nIRCats, however, yielded a reduction in fluorescent response to dopamine in comparison to our standard protocol (*P* = 0.022 significant effect of protocol, two-way ANOVA, Fig. S2). We hypothesize that the dynamic range of the functionalized fibers may have an inherent maximum dictated by the fiber architecture rather than functionalization density. Furthermore, the dynamic range of fluorescent response to dopamine may be attenuated by steric overcrowding of nIRCats immobilized on the fiber surface.

We next considered whether a covalent interaction between SWNT and silica might promote a greater density of dopamine-responsive nIRCats on the fiber surface, as a single nanotube could occupy several amine sites on the silanized silica surface through our standard protocol^53^. Moreover, localized pH changes in a physiological environment could potentially desorb ssDNA from SWNT^54^, thus disrupting the nIRCat-silica bond. A covalent bond, however, is less susceptible pH fluctuation-related cleavage. To this end, we assessed the performance of fibers functionalized with covalently modified SWNTs, which have been shown to maintain analyte-dependent fluorescence responses^55^. We prepared aromatized SWNTs (NH_2_-(GT)_6_-SWNT) as in Chio et al., 2020, then silanized fibers as described followed by the addition of cyanuric chloride to produce triazine-functionalized silica fibers, which were reacted with aromatized nIRCats to covalently link nIRCats to the fibers. This modification yielded fibers with consistent dopamine response but weaker baseline signal (*n* = 6; baseline fluorescence significantly lower than standard protocol, P < 0.001, Wilcoxon ranked-sum; no significant effect of protocol on dopamine response, *P* = 0.317, two-way ANOVA; Fig. S3). We subsequently introduced a 1,3,5-tris(bromomethyl)-benzene (TBMB) reaction, commonly used in the crosslinking of complex polymers^56^, in attempt to produce a silica surface functionalized with reactive bromo-substituted benzylic carbons before reacting the fiber with aromatized SWNTs. This modification yielded fibers with well-quenched fluorescent baselines exhibiting a characteristic SWNT profile (*n* = 5; baseline fluorescence significantly lower than standard protocol, *P* = 0.001, Wilcoxon ranked-sum) and moderate dopamine response, but no improvement of signal compared to our previous electrostatic SWNT-silane manufacturing protocol (no significant effect of protocol on dopamine response, *P* = 0.156, two-way ANOVA; Fig S3).

As nIRCat solution is prepared in water, we sought to assess the effect of solution nucleophilicity on fiber performance by exploring alternative SWNT solution solvents. We desiccated standard aqueous aromatized SWNT solution through rotary evaporation, then attempted to reconstitute the residue separately in four polar aprotic solvents: acetone, dimethylformamide (DMF), dimethyl-sulfoxide (DMSO), and N-methylpyrrolidone (NMP). Reconstitution was successful only with DMSO. Triazine-functionalized fibers incubated in DMSO-SWNT solution yielded baseline fluorescence spectra that were indistinguishable from our standard preparation (*P* = 0.971, Wilcoxon signed-rank) but did not demonstrate the expected fluorescence response to dopamine (significant effect of protocol and significant interaction between protocol and dopamine response, both *P* < 0.001, two-way ANOVA; Fig S3). Together, these experiments provide further insight into the nature of functionalized SWNTs in hydrous and anhydrous environments, in solution and immobilized form.

In summary, we developed catecholamine-sensitive, near-infrared nanosensor-functionalized optical (nIRF) fibers and a compatible dual-nIR mobile fiber photometry imaging platform to readily detect extracellular dopamine *in vitro* and *ex vivo*. We have demonstrated that this tool can be successfully used to detect signals up to 24 weeks post-production and is robust to biofouling anticipated with chronic implantation in biological tissue. This tool introduces the possibility of real-time extracellular catecholamine imaging, potentially permitting a broader range of time-sensitive studies across multiple species with fewer invasive procedures.

In their current form, nIRF fibers can be used as turnkey, shelf-stable, reversible, reusable probes for detection of dopamine at nanomolar levels in solution. We foresee a unique advantage of this tool in the realm of clinical evaluations of catecholamine levels in biofluids such as blood, cerebrospinal fluid, and urine. While blood plasma contains dopamine at sub-nanomolar concentrations^57^, norepinephrine is present at 1.5-1.8 nM^58^. CSF contains dopamine at 1.3-21.7 nM^19,59^ and norephinephrine at up to 25 nM^19,60^, and urine contains dopamine at 65-400 μg/24h^61^ and norepinephrine at 15-80 μg/24h^62^. Elevated plasma and CSF dopamine levels in the 10 nM range are associated with certain pathologies, including coronary artery disease^63^ and psychosis^64,65^. These concentrations are within the dynamic range detectable by nIRF fibers. There is a growing interest in improved biosensors for clinical catecholamine detection from lower-volume samples^66–70^, and we have demonstrated that nIRF fibers can detect dopamine in as little as 10 μL of sample. Moreover, because our sensor platform does not require the addition of subsequent reagents, nIRF fibers permit reversible catecholamine detection in patient samples, which can then be reused for subsequent assays, reducing the overall volume of sample needed in diagnostic testing. Taken together, these attributes elevate nIRF fibers as a promising first-pass diagnostic tool.

Additionally, our work contributes to a greater understanding of SWNT fluorescent dynamics in a unique fiber form factor. We have demonstrated that baseline fluorescence is not necessarily indicative of dopamine detection sensitivity; fibers with low baseline fluorescence perform just as well, if not better than, higher-baseline fibers (Figs. 4, S3). Indeed, prior work has shown that low initial baseline fluorescence of nIRCat dopamine sensors enables larger dopamine-specific responses, and it is possible this phenomenon extends to our nIRF fibers as well^49,71^. Conversely, covalently linking SWNTs to the silica surface of the fiber failed to amplify dopamine response, which we postulate could be due to unintended reactions between the halide-functionalized silica and nucleotides in the ssDNA oligomer or pH-driven desorption.

One limitation of this tool is the apparent attenuation of fluorescence observed after long-term exposure to brain homogenate. This could be due to biofouling compromising the exciton recombination efficiency of nIRCats or endonucleases compromising the integrity of the (GT)_6_ functionalization on nIRCat surfaces. Biofouling may be mitigated by passivation of SWNTs with polyethylene glycol, which reduces SWNT-induced platelet aggregation^72^, permits circulation *in vivo*^43^, improves targeted drug delivery success^73^, minimizes cytotoxicity and nonspecific cellular interactions^74^, reduces expression of inflammatory cytokines, attenuates microglial activation, and further enhances dopamine imaging^50^. An alternative approach to nIRF fiber synthesis could encapsulate nIRCats in a hydrogel, which has demonstrated excellent biocompatibility and light-guiding properties^75^. This strategy has successfully enabled SWNT-based sensors for detection of steroid hormones^76^, essential vitamins^77^, and chemotherapeutic drugs^78^ *in vivo*. Moreover, this strategy has recently proved applicable to large mammal models for multi-week sensing without adverse health effects^79^, suggesting tractability across species. Further investigation combining these strategies may present a viable route by which to successfully utilize nIRCats in living brain tissue.

Overall, we foresee this tool as a starting point from which to expand resources available for quickly assessing biofluid catecholamine levels, chronic catecholamine monitoring in genetically intractable or time-sensitive model systems, or and study of the effects of catecholamine pharmacology. These data represent the first step in development of an optical probe for catecholamine detection that may provide a real-time readout of catecholamine release during human surgery or the ability to chronically monitor catecholamine levels in a patient over time, leading to a deeper understanding of the catecholamine dynamics behind the progression and treatment of debilitating human illnesses.

## Author Contributions

Project was conceived by M.P.L. and L.W. All authors contributed to experiment design. M.E.K. performed all experiments except protocol modification experiments, which were performed by R.A.M. M.E.K. analyzed all data. M.E.K. wrote most of the manuscript with contributions from R.A.M, M.P.L and L.W.; all authors edited and approved the final manuscript.

## Notes

**The authors declare no competing financial interest**.

## ACKNOWLEDGMENTS

We thank J. Travis Del Bonis-O’Donnell, PhD, for feedback and direction regarding silanization and M. Moein Safaee, PhD, for assistance constructing the fiber photometry rig.

