## Supplementary Information for "Optical Fibers Functionalized with Single-Walled Carbon Nanotubes for Flexible Fluorescent Catecholamine Detection"

### Materials and Methods

Fiber preparation (standard):
Aminosilanes are a class of compounds commonly used as coupling agents for silica-based materials such as glass. In solution, the amine groups interact with negatively charged molecules, such as DNA. We leveraged this interaction to develop an aminosilane-based protocol, loosely based on previous demonstrations of silica silanization (Zhu, Lerum, & Chen, 2012), to immobilize nIRCats on glass optical fibers for photometric measurement of dopamine. We used a 400µm-diameter core multimode silica optical fiber (Thorlabs FP400URT, 0.5 NA) glued with epoxy into a 1.25mm diameter iron ferrule (Thorlabs SFLC440). The exposed length of fiber was trimmed to 4.5mm, the approximate depth of the rodent nucleus accumbens from the brain surface. Fibers were first cleaned by immersion for at least five minutes each in acetone, isopropyl alcohol, and molecular grade water, in that order.
*Hydroxylation of silica surface*: This procedure was incorporated only for data represented in figure 3. Fibers were soaked in potassium permanganate solution for 30 minutes, washed repeatedly with water, soaked in a solution of potassium hydroxide in ethanol for 30 minutes, again washed with water, then placed in a glass vial which had been pre-treated with piranha solution (1:1 (v/v) concentrated sulfuric acid and 30% hydrogen peroxide). 300µL fresh piranha solution was added to the vial and fibers were incubated for 30 minutes. Fibers were then washed with water to remove ionic debris, ethanol to remove water and less polar contaminants, and toluene to remove ethanol.
*Silanization*: Prepared fibers were placed in an oven-dried 3-necked round-bottom flask containing 19ml anhydrous toluene. The flask was connected to a condenser and continuously flushed with nitrogen gas. Exposed flask necks were secured with septa, and 1ml (3-Aminopropyl)triethoxysilane (APTES, 99%, Sigma-Aldrich) was injected through one neck to create a 5% APTES solution in toluene. To increase APTES-silica hydrolyses and minimize APTES polymerization, we aimed to attenuate atmospheric humidity. The flask was then lowered into an 80°C heated oil bath and the condenser was flushed with cold tap water to reduce evaporate throughout the one-hour silanization reaction.

*Functionalization with nIRCats*: Post-silanization, tips were immersed for at least five minutes each in toluene, ethanol, and molecular grade water, in that order, to displace any residual weakly bonded silanes. Fibers were then dried overnight in a 110°C oven to promote formation of siloxane bonds. Afterwards, fibers were incubated in 50-200 mg/L nIRCats solution (manufactured in-house as in Beyene et al., 2019) for 30-60 minutes. Fibers were passively incubated or incubated with bath sonication for 30 minutes followed by a rest of at least 30 minutes. Immediately after nIRCats incubation, fibers were transferred to 1x phosphate buffered saline (PBS, Gibco) for at least ten minutes before spectra were assessed. Functionalized fibers can be stored long-term in clean plastic or glassware.

Production of covalently-modified nIRCats: SWNTs subjected to an aromatization reaction using cyanuric chloride and sodium azide can be solubilized in water via functionalization with ssDNA sequences without substantial loss of analyte-dependent fluorescence response (Chio et al., 2020). We adapted the protocol described in Chio et al., 2020, to produce covalently-modified nIRCats. In brief, pristine SWNTs were reacted with cyanuric chloride and sodium azide to produce TrZ-L-SWNT, which were subsequently reacted with ethylenediamine to produce $NH_2$-SWNT. $(GT)_6$-ssDNA was adsorbed to $NH_2$-SWNT as in Beyene et al., 2019, to yield $NH_2$-$(GT)_6$-SWNT, which was evaluated for DA response.

Covalent nIRCats-silica linkage:
*Triazine functionalization*: Fibers were washed, etched, hydroxylated, and silanized as described above, then reacted with 1 g cyanuric chloride in 20 mL N-methyl-2-pyrrolidone (NMP) at 70°C for 24 hours to produce triazine-functionalized silica fibers. Triazine-functionalized fibers were then incubated in an aqueous solution of $NH_2$-$(GT)_6$-SWNT for 24 hours.

*1,3,5-tris-(bromomethyl)benzene functionalization*: Fibers were washed, etched, hydroxylated, and silanized as described above, then reacted with 20mL dimethylformamide, 250 µL triethylamine, and 250 mg TBMB at 70°C for 24 hours under nitrogen. Fibers were then incubated in a solution of $NH_2$-(GT)$_6$-SWNT for 24 hours.

Exchange of SWNT solvents:
*Chemical desiccation:* In brief, nIRCats prepared as in Beyene et al., 2019 or $NH_2$-(GT)$_6$-SWNT as described above were combined with a water-miscible polar aprotic solvent, then a desiccant was added to remove water. 50 µL each of acetone, dimethylformamide (DMF), dimethylsulfoxide (DMSO), and NMP, and were added to separate aliquots of 50 µL nIRCat solution, then agitated. 50 µL solvent was subsequently added with agitation between each addition until the total volume of solution reached 250 µL. Approximately 100 mg anhydrous sodium sulfate was added and solutions were stored at 10°C.
*Evaporation and reconstitution:* nIRCats prepared as in Beyene et al., 2019 or $NH_2$-(GT)$_6$-SWNT as described above were evaporated under reduced pressure, then the resulting residue was vigorously agitated with water or with polar aprotic solvents. 250 µL aqueous SWNT solution was added to a 50 mL round bottom flask and the water was removed under reduced pressure using a rotary evaporator. 250 µL of water, NMP, or DMSO was added, and the residue was agitated under the solvent. The solvent was collected and centrifuged at $2x10^4$ rcf for 30 minutes at room temperature, and the supernatant was collected and characterized by UV-Vis absorbance and IR fluorescence spectroscopy. An approximate extinction coefficient for SWNT in DMSO at 632 nm was calculated by serial dilution of known-concentration aqueous $NH_2$-(GT)$_6$-SWNT in DMSO followed by averaging the calculated values. This value was then used to approximate the concentration of SWNT in DMSO.

Laser/spectrometer data collection: A 721-nm laser (Opto Engine LLC) was fiber-coupled to an inverted Zeiss microscope (Axio Observer D1). A 60cm-long sheathed fiber with a plastic ferrule at one end was coupled directly to the light source in place of an objective and provided a conduit for excitation light. Each individual nIRF fiber was then plugged into the plastic ferrule at the end of the conduit fiber; laser power exiting the fiber was measured to be 4-8 mW. Each nIRF fiber was suspended in 40 ml 1x PBS. Fluorescence spectra were collected from 850-1300 nm by a Princeton Instruments spectrograph (SCT 320) and a liquid nitrogen–cooled Princeton Instruments InGaAs linear array detector (PyLoN-IR). Each fiber was equilibrated to the solution and laser light for ten minutes in 1x PBS with the microscope shutter open before baseline spectrum collection. Thereafter, a 1 mM DA solution was added directly into the PBS in increments of 0.4 µl, 4 µl, 40 µl, and 400 µl to produce 10 nM, 100 nM, 1 µM, and 10 µM DA solutions, respectively, unless otherwise noted. Spectra were collected one minute after each subsequent addition of DA. When assessing spectra in small volumes, fibers were suspended in individual 10 µl aliquots of 10 nM, 100 nM, and 1 µM DA. Fibers were equilibrated for 10 minutes in each solution before data was acquired.

Solutions:
*Artificial cerebrospinal fluid (aCSF):* aCSF was prepared in-house, consisting of sodium chloride (6.90 g/L), potassium chloride (0.26 g/L), hydrated magnesium chloride (0.264 g/L), sodium phosphate (0.12 g/L), sodium bicarbonate (2.20 g/L), D-glucose (1.98 g/L), and hydrated calcium chloride (0.37 g/L) in one liter of MilliQ water.
*Brain homogenate:* the brain of one adult male mouse was extracted in accordance with laboratory animal care guidelines. 5 mL 1x PBS was immediately added to the brain and homogenate was generated by probe-tip sonication for 30 minutes.
*Blood plasma*: frozen 1 mL aliquots of human blood plasma (pooled, Lee Biosciences) were thawed and diluted with 1x PBS.

Dual-nIR fiber photometry rig and electrical stimulation: We designed and constructed a mobile dual-nIR fiber photometry rig for use with nIRF fibers in *ex vivo* and *in vivo* applications. A 635 nm excitation laser ( 35-50 mW at end of conduit fiber) was fiber coupled, collimated, and deflected into a patch fiber by a 900nm long-pass dichroic mirror. The patch fiber allows for individual nIRF fibers to be implanted in tissue and connected when needed via a plastic sleeve. Emission light from nIRF fibers travels through the same patch fiber and passes through the dichroic mirror to an ultrasensitive, thermoregulated nIR camera (Ninox 640 II, Raptor Photonics). The patch fiber was affixed to the camera lens such that the only light entering the lens was from the fiber itself. For each acquisition, Micro-Manager imaging software (v. 1.4.23) acquired 600 frames at 8.33 Hz with 0.5 mA electrical stimulation triggered at frame 200. All nIRF fibers used on this rig are first evaluated for *in vitro* DA response with the spectrometer.

Analysis of fluorescence: For spectroscopic data, we integrated fluorescence values (arbitrary units) from 1050-1300 nm for all readings. Each fiber's fluorescence in 1x PBS served as the baseline against which subsequent measurements from that fiber were normalized. For nIR camera images, image acquisition was controlled by Micro-Manager software, which summed pixel intensity from a 640 x 512 pixel field of view, an area which encapsulated the totality of the 400 μm-diameter nIRF fiber surface. A line fit between the average of the first 50 frames and average of the last 50 frames of each acquisition was taken as the fluorescence baseline to correct for drift; all fluorescence values are represented normalized relative to that baseline. All analysis was performed in MATLAB using custom scripts.

**Supplementary Discussion:**

We recognize that several methods currently exist to measure catecholamine levels in human and animal subjects, though each have caveats. Magnetic resonance imaging (MRI) can detect tissue iron, a proxy measurement of dopamine receptor density used to infer the development of catecholamine-releasing terminals, but cannot report real-time release dynamics or short-term plasticity[38–40]. Positron emission tomography (PET) imaging can quantify catecholamine transmission but requires injection of a radioactive receptor-binding ligand into a patient's vein, and the detected ligand signal cannot be separated from its metabolite[37]. Probe implantation for microdialysis[41,42], or fast-scan cyclic voltammetry (FSCV)[43,44] permits direct analysis of catecholamine release, but it is highly invasive, and FSCV, while more precise, has not yet been implemented for regular clinical use due to safety and standardization concerns[45–47]. A less invasive strategy is to monitor catecholamine levels in bodily fluids, such as patient cerebrospinal fluid, blood serum, saliva, or urine samples. Altered catecholamine levels in biofluids can be an early marker of disease, including neuroblastoma[48–50], Parkinson's Disease[51–54], Alzheimer's Disease[20,55,56], and psychosis[57,58]. Conventionally, catecholamines are quantified using high-performance liquid chromatography with tandem mass spectrometry or electrochemical detection[59,60]. In such assays, aliquots of sample are added to reagents, then run through a multi-step process, which is irreversible, expensive, and time-consuming. In combination with concurrent laboratory tests, dozens of milliliters of fluid may need to be collected from venous or lumbar puncture, which can contribute to patient suffering.

In animal research, fluorescent sensors from modified catecholamine receptors have been developed in the last decade that enable optical measurement of catecholamine dynamics[64,65]. These have been heavily reviewed elsewhere[37,66]. While these tools have revolutionized *in vivo* preclinical studies, they require viral transfection or genetic modification for protein expression, which limits their translatability to other model organisms or clinical applications. They are also incompatible with pharmacology as dugs intended to act on endogenous receptors also bind to these reporter tools.

We also recognize that one barrier to *in vivo* use of our nIRF fiber is that the long-term effects of SWNT exposure are not fully known[110–113]. While our work has shown no measurable SWNT desorption from functionalized nIRF fibers, solution-phase SWNT can induce adverse biological effects at certain concentrations and exposure durations. For instance, there is some evidence that chemically functionalized multi-walled carbon nanotubes induce a transient immune response in the brain and that SWNTs alter microglial morphology in cell culture[80,114,115], but it remains unclear how SWNTs affect the brain *in vivo*. As such, future research should take care to more closely investigate the potential toxicity of single-walled carbon nanotubes in brain tissue, and whether their immobilized form function mitigates these toxicities. Given the lack of evidence of SWNT desorption from the fiber, we anticipate this concern may be mitigated by combination with a functionalized fiber.

To our knowledge, however, there exist few methods by which to safely and continuously measure catecholamines in the human brain or biofluids without the limitations of microdialysis, FCSV and GEDIs indicated above. The nIRF fiber may be able to mitigate risk of use by reducing size, removing need for electrical stimulation, and reducing need for use of viral transfection or other irreversible treatment (Fig. S4). The small size of the optical fiber may permit relatively easy implantation in live brains, and compatibility with pharmacology means these fibers could be used alongside ongoing clinical trials of new drugs that target dopamine pathways to measure changes in dopamine release without signal interference from the drug's mechanism of action. Furthermore, growing interest in translational fluorescent sensors indicates a future rise in infrastructural support for this technology[116].

**Supplementary Figures:**

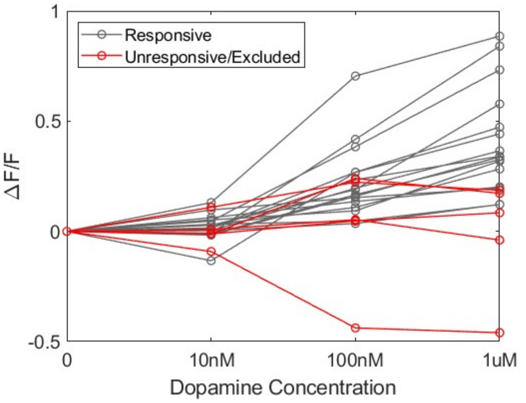

**Figure S1.** Integrated fluorescence from 1050-1300nm for increasing concentrations of dopamine for individual fibers. Gray = responsive fibers (in Fig 2A) vs. red = unresponsive fibers excluded from analysis.

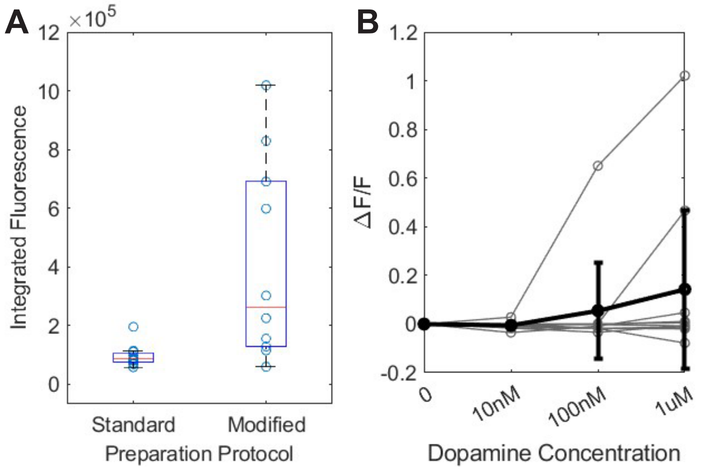

**Figure S2**. A) Protocol modifications significantly increased baseline fluorescence (standard protocol, n = 16; modified protocol, n = 10). B) Modifications yielded a reduced response to dopamine with 7/10 fibers exhibiting no response.

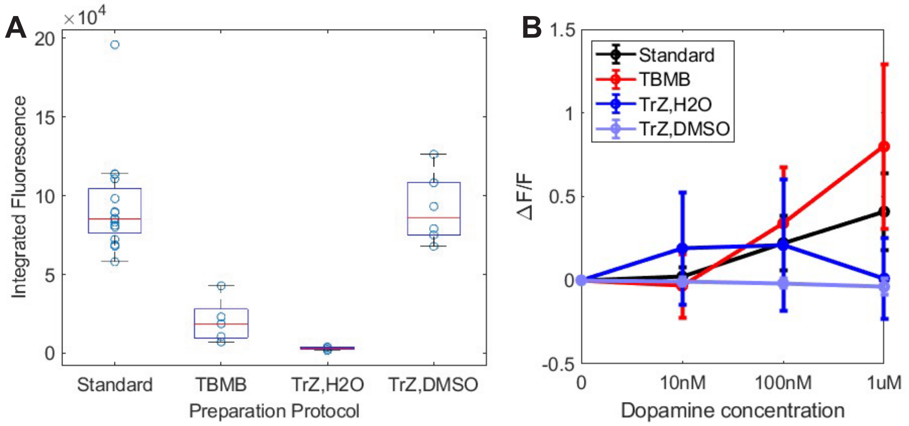

**Figure S3.** A) Baseline fluorescence and B) Dopamine response profiles of alternative nIRF fiber manufacturing protocols.

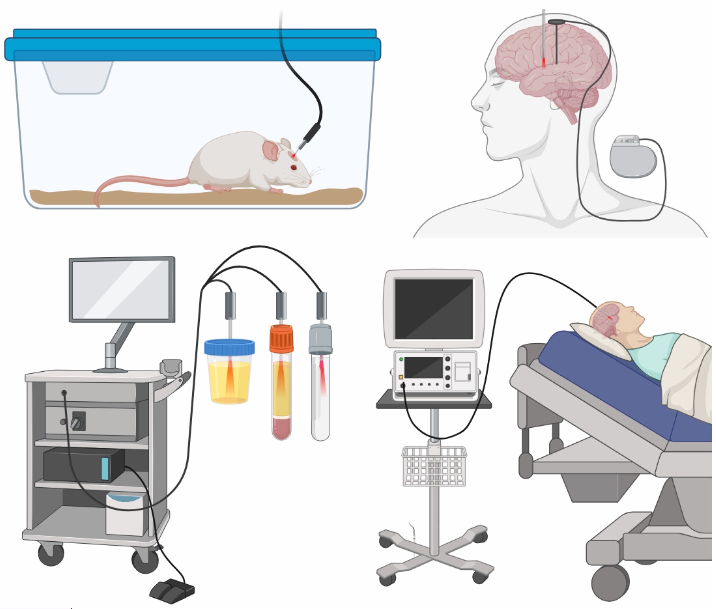

**Figure S4. Potential applications of the nIRF fiber.** We envision this tool providing utility for (clockwise from top left) behavioral neuroscience experiments in animal models, chronic implantation for monitoring patient catecholamine release with concurrent deep-brain stimulation, acute implantation for monitoring patient catecholamine release during brain

**Commented [ML1]:** Uncommon to have a figure without data. Recommend combining fig 1 and 2

**Commented [MK2R1]:** This figure was Linda's suggestion as a summary - I'm happy to couple it to Fig 2, but I'm not sure it makes sense with the qualitative data shown there. Given the scope of the paper, I'd propose simply removing it - I'll give you final say

**Commented [ML3R1]:** Agree on removing from main paper - can you please place it in the SI? Be sure to renumber figures there accordingly.

surgery, and preliminary analysis of catecholamine levels in clinical samples of biofluids. Figure created with BioRender.com.

**Table S1**. Summarized experiment responses to dopamine ($\Delta F/F$, mean ± SD)

| **Solvent** | 40mL 1x PBS | 40mL 1x PBS | 40mL 1x PBS | 40mL 1x PBS | 40mL 1x PBS | 40mL diluted blood plasma | 40mL aCSF | 10μL aCSF | 10μL undiluted blood plasma |
|---|---|---|---|---|---|---|---|---|---|
| **Time of measure-ment** | Mfg. date | Post incubation in blood plasma | Post incubation in brain homog. | After 24 days stored in 1x PBS | After 24 weeks stored in air | After 24 weeks stored in air | After 24 weeks stored in air | After 24 weeks stored in air | After 24 weeks stored in air |
| ***n*** | 16 | 6 | 4 | 6 | 4 | 4 | 4 | 4 | 1 |
| **10nM dopamine** | 0.022 ± 0.056 | 0.033 ± 0.025 | -0.004 ± 0.053 | 0.050 ± 0.034 | 0.024 ± 0.053 | 0.103 ± 0.086 | 0.063 ± 0.048 | 0.114 ± 0.069 | 0.182 |
| **100nM dopamine** | 0.223 ± 0.162 | 0.198 ± 0.118 | 0.098 ± 0.050 | 0.334 ± 0.156 | 0.266 ± 0.174 | 0.307 ± 0.143 | 0.190 ± 0.106 | 0.118 ± 0.043 | 0.221 |
| **1μM dopamine** | 0.411 ± 0.230 | 0.381 ± 0.189 | 0.278 ± 0.073 | 0.790 ± 0.376 | 0.517 ± 0.235 | 0.467 ± 0.152 | 0.289 ± 0.126 | 0.143 ± 0.035 | 0.856 |

Commented [ML4]: Suggest moving to SI since doesn't present any new data

**Table S2.** Results of modifications to standard nIRF fiber preparation protocol.

| Anhydrous SWNT solution | | |
|---|---|---|
| Solution | Concerns | |
| Acetone | Immediately precipitated | |
| NMP | Uncertain extinction coefficient; failed reconstitution | |
| DMF | Overlapping emission spectra | |
| DMSO | None | |
| SWNT-glass covalent attachment | | |
| Coupling Agent | Immobilization on fiber | DA response from fiber |
| TrZ-functionalized glass + NH2-(GT)6-SWCNT in water | successful | No significant improvement over non-covalent protocol |
| TrZ-functionalized glass + NH2-(GT)6-SWCNT in DMSO | successful | No DA response |
| TBMB + NH2-(GT)6-SWCNT in water | successful | No significant improvement over non-covalent protocol |
| TBMB + NH2–(GT)6–SWCNT in DMSO | unsuccessful | N/A |
| TBMB + NH2–(GT)6–SWCNT in NMP | unsuccessful | N/A |
